# BWA-MEME: BWA-MEM emulated with a machine learning approach

**DOI:** 10.1101/2021.09.01.457579

**Authors:** Youngmok Jung, Dongsu Han

## Abstract

**Motivation:** The growing use of next-generation sequencing and enlarged sequencing throughput require efficient short-read alignment, where seeding is one of the major performance bottlenecks. The key challenge in the seeding phase is searching for exact matches of substrings of short reads in the reference DNA sequence. Existing algorithms, however, present limitations in performance due to their frequent memory accesses.

**Results:** This paper presents BWA-MEME, the first full-fledged short read alignment software that leverages learned indices for solving the exact match search problem for efficient seeding. BWA-MEME is a practical and efficient seeding algorithm based on a suffix array search algorithm that solves the challenges in utilizing learned indices for SMEM search which is extensively used in the seeding phase. Our evaluation shows that BWA-MEME achieves up to 3.45x speedup in seeding throughput over BWA-MEM2 by reducing the number of instructions by 4.60x, memory accesses by 8.77x, and LLC misses by 2.21x, while ensuring the identical SAM output to BWA-MEM2.

**Availability:** The source code and test scripts are available for academic use at https://github.com/kaist-ina/BWA-MEME/.

## 1 Introduction

DNA sequencing has become a critical piece in modern medicine, advancing the practice in disease diagnosis, prognosis, and therapeutic decisions. The state-of-the-art DNA sequencing method is called next-generation sequencing (NGS). Modern NGS hardware generates billions of short reads in a single run. This, in turn, requires the alignment of short reads (i.e., short DNA fragments) to the reference DNA sequence. As large-scale DNA sequencing operations run hundreds of NGS, developing an efficient short read alignment algorithm has become ever more important.

The state-of-the-art alignment process is divided into two phases, seeding and extending, following the seed-and-extend paradigm (Li, 2013; Li and Homer, 2010; Liu and Schmidt, 2012; Liu et al., 2012). The seeding phase searches for exact matches of seeds (substrings of short reads) in the reference DNA sequence, which identifies the possible alignment positions of the short read in the reference DNA sequence. In the extending phase, the seeds from the earlier phase are extended. In the process, the alignment scores of the extended seeds are calculated using the Banded-Smith-Waterman (BSW) algorithm (Vasimuddin et al., 2019). Many studies (Ahmed et al., 2015; Ho et al., 2019; Houtgast et al., 2018; Subramaniyan et al., 2021; Tárraga et al., 2014) have shown that the seeding phase is a major performance bottleneck in the popular alignment software BWA-MEM2 (Vasimuddin et al., 2019) and Bowtie 2 (Langmead and Salzberg, 2012). In particular, finding an exact match of short reads— or substrings of short reads—within the reference DNA sequence, is the main problem that fundamentally constrains the performance of the seeding phase (Langmead and Salzberg, 2012; Li, 2013; Li et al., 2009). Thus, in this paper, we focus on the seeding phase which involves only the exact matching.

For efficient exact match search, it is necessary to index the reference DNA sequence and perform an in-memory index lookup. Recently, several new index structures have been studied to solve the exact match search problem in an efficient manner. The state-of-the-art index structures fall into two major categories: traditional index-based (Subramaniyan et al., 2021; Vasimuddin et al., 2019) and machine-learning-based (Ho et al., 2019, 2021; Kirsche et al., 2021). Examples of the traditional index structures are FM-index (Langmead and Salzberg, 2012; Li, 2013; Li and Durbin, 2009; Li et al., 2009) and enumerated radix tree (ERT) (Subramaniyan et al., 2021) index. FM-index is a compressed version of the suffix array (Ferragina and Manzini, 2001), which progressively extends a substring from a single character and finds its exact match in the reference DNA sequence. We refer to the substring of the short read that is given as input to the exact match search problem as a substring. The typical length of a short read is between 100 to 300. To speed up the process, the ERT index uses an enumerated index table for finding the exact match of the first 15 consecutive bases of the substring. After this, ERT utilizes a radix tree that encodes the suffixes of the reference DNA sequence, which naturally supports multi-character lookups. Despite this, both FM-index and ERT index requires *O*(*N*) memory accesses, where N is the length of the input substring.

LISA (Ho et al., 2019, 2021) and Sapling (Kirsche et al., 2021) use machine-learning-based index structures. LISA proposes a new data-structure called IP-BWT. IP-BWT employs a learned index that supports an exact match lookup in the suffix array. Although it is still a linear time algorithm, using IP-BWT requires fewer memory accesses compared to the original FM-index because it matches 21 bases in a single lookup. However, LISA assumes seed search only starts at the first base of the short read, whereas the seeding algorithm in BWA-MEM2 requires to start the search at an arbitrary point in the short read. Thus, LISA cannot be used as a replacement of BWA-MEM2. Sapling (Kirsche et al., 2021) has shown that employing a learned index in suffix array search can outperform FM-index based algorithms. However, Sapling only works for input substrings that have the perfect match to the reference DNA sequence. Therefore, it cannot be used for real-world alignment where the input may not have a perfect match to the reference DNA sequence.

The key limitation of existing index structures is that it requires memory accesses and instructions proportional to the length of the substring. In reality, seeds are of variable size, and thus the index structure must support a fast exact match search for arbitrary length substrings. Our central tenet is that to obtain high seeding throughput, we must minimize the number of memory accesses required for exact match search of arbitrary length substrings and break the strong dependency between the length of substring and the number of memory accesses.

In this paper, we present BWA-MEME, the first alignment software that performs exact match search with *O*(1) memory accesses leveraging the learned index. Specifically, we design the first full-fledged learned-index-based seeding algorithm that achieves constant lookup time for an exact match search of arbitrary length substrings. BWA-MEME significantly reduces the number of instructions, memory accesses, and LLC misses, in particular, for the long substrings.

Building a full-fledged alignment software that leverages learned index in suffix array search, however, involves solving a number of new and non-trivial challenges. First, the learned index has to provide an accurate exact match position of an arbitrary substring in the suffix array. However, it is difficult to guarantee high prediction accuracy in the suffix array which is large and has an imbalanced distribution of suffixes. Furthermore, variable-length suffix or substring must be encoded into the numerical key to use the learned index. Second, using the suffix array search requires a new design for the super-maximal exact match (SMEM) search (Li, 2012, 2013). SMEM search algorithm must find all seeds covering the given point of the short read, while minimizing the number of exact match searches. Seeding algorithm uses a hit threshold to find seeds that have multiple hits in the reference DNA sequence. Thus, the SMEM search algorithm is required to find seeds with the maximum number of hits but is less than or equal to the hit threshold. Finally, minimizing the memory accesses and CPU cache misses introduced by using a learned index is important.

BWA-MEME addresses these challenges by introducing a new learned index structure and algorithms. First, we present a partially-3-layer recursive model index (P-RMI) which adapts well to the imbalanced distribution of suffixes and provides accurate prediction. In this process, we design an algorithm that encodes the input substring or suffixes into a numerical key. The numerical key is given as input to P-RMI where P-RMI provides predicted position in the suffix array and error bound for the prediction. Binary search is performed within the error bound to find the reference position where the substring aligns to. Second, we present an efficient SMEM search algorithm that uses the same or less number of exact match searches compared to the state-of-the-art SMEM search algorithms. Finally, we reuse the lookup result to exploit the redundancy of the input substrings to the exact match search problem. This further reduces the number of memory accesses and CPU cache misses that occur during the seeding phase.

Our evaluation shows that, 1) BWA-MEME achieve up to 3.45x and 1.42x speedups in seeding throughput and alignment throughput respectively over BWA-MEM2, while ensuring identical output; 2) BWA-MEME drastically reduces the number of instructions executed by 4.60x, memory accesses by 8.77x, last-level-cache (LLC) misses by 2.21x, and data fetched per read by 4.41x compared to seeding algorithm of BWA-MEM2; and 3) BWA-MEME provides options to balance the trade-off between alignment throughput and the required memory space.

## 2 Background

### 2.1 The Seeding algorithm of BWA-MEM2 and ERT

A maximal-exact match is a substring (of short reads) that cannot be further extended in either direction without a change in the number of hits (exact matches) to the reference DNA sequence. A SMEM is a unique MEM that is not contained in other MEMs. To find all positions of seeds where the short read may potentially align, the seeding algorithm executes a super-maximal exact match (SMEM) search multiple times with various pivot points, minimum seed length thresholds, and hit thresholds. SMEMs found in the SMEM search that are longer than the minimum seed length threshold are selected as seeds. In the following, we denote the SMEM search algorithm of BWA-MEM2 (Vasimuddin et al., 2019) and ERT (Subramaniyan et al., 2021) as SMEM-BWA and SMEM-ERT, respectively.

#### SMEM search algorithm and extension

We first describe the extension used in SMEM search and explain how extensions are performed to find SMEMs. Let *S* be a short read, and *S*[*i, j*] denote the substring between position *i* and position *j* of the short read. Extension from position *P_s_* is finding farthest position *P_e_* where the substring *S*[*P_s_, P_e_*] has a maximum number of hits but is less than or equal to the hit threshold. Therefore, for each extension, the exact match search algorithm is used to find the number of hits for substring *S*[*P_s_, P_e_*]. Depending on the direction of extension, it is called forward extension if *P_s_ < P_e_* and otherwise backward extension. We denote each point where forward and backward extension end as *forward*(*p*) and *backward*(*p*), respectively.

The goal of an SMEM search algorithm is to find all SMEMs that include the pivot point *P_pivot_*. As SMEMs are substrings that cannot be further extended, all SMEMs that include *P_pivot_* can be found by; 1) performing a forward extension from *P_pivot_* of the short read and; 2) performing a backward extension from every points in [*P_pivot_, forward*(*P_pivot_*)]. This finds all SMEMs that include *P_pivot_* but incurs excessive computation.

#### SMEM search algorithm of BWA-MEM2

Instead of performing backward extension in all points in [*P_pivot_, forward*(*P_pivot_*)], SMEM-BWA performs backward extension only in the point *p* where substring *S*[*P_pivot_, p*] and substring *S*[*P_pivot_, p* + 1] have different number of hits. This is because for ∀*p* ∈ [*P_pivot_, forward*(*P_pivot_*)], if substring *S*[*P_pivot_, p*] and substring *S*[*P_pivot_, p* + 1] have the same number of hits, *backward*(*p*) and *backward*(*p*+1) are identical. Therefore, during the forward extension from *P_pivot_*, all positions where a number of hits changes are marked as left extension points (LEPs).

Figure 1 illustrates the design. **(1) Determining LEPs:** SMEM-BWA starts with the forward extension of a single character at the pivot point of the short read. During the forward extension, substrings are extended one character at a time using the FM-index. The LEP bit is set to 1 when the number of hits changes from the preceding substring. In Figure 1, the number of hits decreases as the length of forward extended substring increases, and the LEP bit is set to 1 when the number of hits changes. Each dashed line box represents a single extension. **(2) SMEM search:** The backward extension is performed in the positions where the LEP bit is set to 1. After the backward extension is performed in each position, SMEMs are selected from the backward extended substrings. SMEMs are backward-extended substrings that are not contained in other substrings and are longer than the minimum seed length threshold. In Figure 1 backward extension is not performed in the substrings with LEP bit set to 0 which is labeled “Not Extended”. Substrings that are contained in other longer substrings are labeled “Contained", and substrings that are not contained in the other substrings are selected as SMEM which are labeled “SMEM”.

**Fig. 1.**
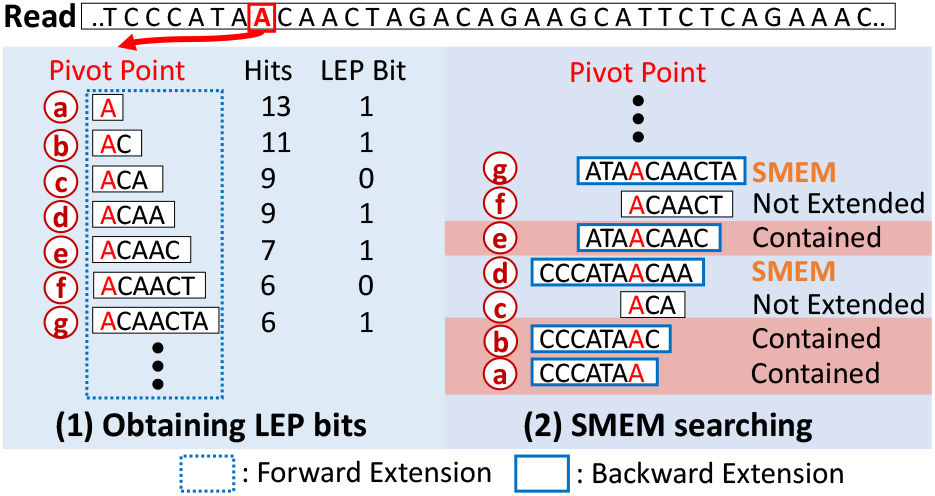
SMEM Search Algorithm of BWA-MEM2 (SMEM-BWA algorithm)

#### SMEM search algorithm of ERT

One limitation of SMEM-BWA is that it still performs backward extensions on substrings that eventually do not become SMEM. To overcome this limitation, SMEM-ERT performs extensions in a zigzag fashion. Similar to SMEM-BWA, SMEM-ERT performs backward and forward extension in positions where the LEP bit is set to 1. This is intended to avoid finding duplicate SMEMs and reduce redundant extensions.

Figure 2 illustrates the two stages of SMEM-ERT: **(1) Obtaining LEP bits:** SMEM-ERT starts with a forward extension at the pivot point of the short read. LEP bits for all characters in the forward extension are obtained from the ERT index. **(2) SMEM searching in zigzag fashion:** Starting at the pivot point, the backward and forward extensions are repeatedly performed until the forward extension reaches the end of the obtained LEP bits. The forward extension starts at the point where the backward extension ends, and the backward extension always starts at the nearest point where the LEP bit is set to 1, as illustrated in Figure 2.

**Fig. 2.**
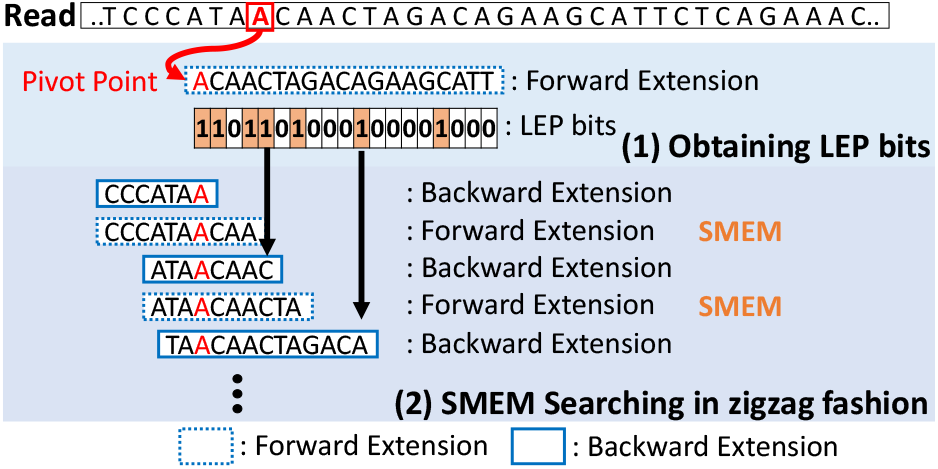
SMEM Search Algorithm of ERT (SMEM-ERT algorithm)

**Fig. 3.**
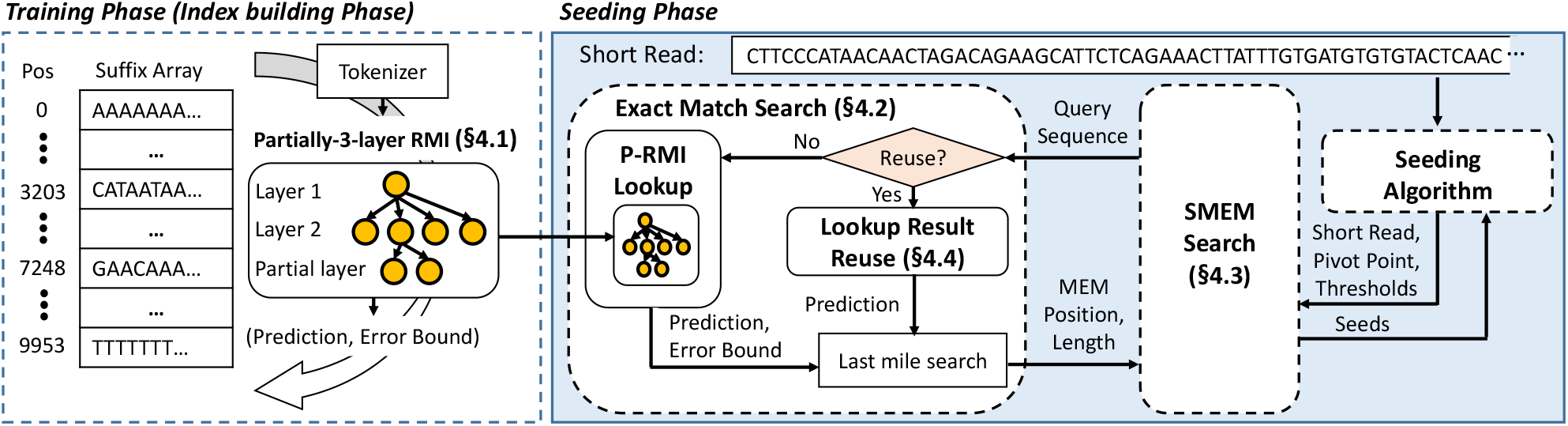
BWA-MEME: Design Overview

### 2.2 Recursive Model Index Structure

Learned indices use machine learning (ML) models (e.g. linear regression model) to replace traditional index structures or search methods used for sorted key-value data (e.g. B-tree, binary search). Recursive model index (RMI) (Kraska et al., 2018) is a commonly used hierarchical structure for learned indices, which uses multiple layers of ML models. The first layer usually has a single model and subsequent layers have a progressively larger number of models. The algorithm of RMI consists of the training and lookup phase. For simplicity, we refer to a ML model as a model.

#### Training phase

Training is performed layer by layer starting from the first layer. Model in the first layer is trained with whole key-value data to predict the position of keys accurately. Then keys are distributed to the models in the second layer according to the predicted position of keys. Models in the second layer are trained with the assigned key-value data and keys are again distributed to the models in the third layer. This process continues until the models in the last layer are trained. As prediction may have an error, RMI guarantees a search bound for keys seen in the training phase. The error bound of leaf models is calculated by iterating through keys assigned to each leaf model. We refer to models that are used for the output of RMI as the leaf models. Each leaf model stores the calculated error bound to be used later in the lookup phase.

#### Lookup phase

The lookup phase of RMI consists of two stages. **Stage 1) Prediction**: Given a key, the first layer model predicts the position of the key, and one of the models in the second layer is selected according to the prediction. The selected model in the second layer subsequently makes a prediction, and one of the models in the third layer is selected. RMI repeats this process recursively until the model in the last layer is selected. The prediction made in the last layer is used as the output of RMI. **Stage 2) Last mile search**:The true position of the key is searched starting from the predicted position of RMI. Binary search can be used if an error bound for prediction is defined in the training phase. Otherwise, linear search or exponential search is used.

## 3 Motivation and Goal

To solve the exact match search problem with minimal time, it is necessary to index the reference DNA sequence and perform an in-memory index lookup. However, it is a challenging task to build an index that fits in limited memory and supports exact match search of substring (which have a length between 0 to 300) in a long reference DNA sequence (e.g. 3 billion lengths for human reference DNA).

### Why use learned index?

Applying machine learning based indexing in the short read alignment is attractive considering that reference DNA sequence does not change frequently. Once the models are trained, they can be used without further training unless the new version of the reference DNA sequence is made. In addition, most of the short reads have the perfect match to the reference DNA sequence (Consortium et al., 2015), which means that the test data and the train data are similar. This makes the machine-learning based approach even more effective.

### Why apply learned index on suffix array?

A suffix array is an array that stores the positions of suffixes sorted in lexicographical order. For a human reference DNA sequence whose size is 6G bases (including its reverse complement), the corresponding suffix array is 31 GB. However, suffix array search has not been used for the exact match search problem because it is slow when combined with the binary search which requires *O*(*log*(|*R*|)) times of memory accesses. Therefore, FM-index applies Burrows-Wheeler transform (BWT) (Li and Durbin, 2009) on suffix array, which reduces the number of memory accesses from *O*(*log*(|*R*|) to *O*(*N*), where N is the length of the substring. Recently, LISA presented IP-BWT which integrates learned index to FM-index. In particular, LISA processes 21 bases in a single learned index lookup followed by additional binary search in IP-BWT. For the input substring that is longer than 21, LISA requires multiple learned index and IP-BWT lookups, which results in memory accesses proportional to the length of substring. Therefore, the number of memory accesses in LISA is lower bounded by the length of substring, even if the learned index predicts the exact position.

In contrast, directly employing a learned index to suffix array search without using a compressed structure of FM-index, requires number of memory accesses independent to the length of input substring. This is because the number of memory accesses required in the suffix array search is bounded by the errors of the leaf models in P-RMI which is a constant value determined at the training phase. The learned index itself requires a single memory access to find the exact match in the best case, which is the minimum achievable value for the exact match search problem. Hence, to achieve the minimal lookup time, we choose to utilize learned index on the suffix array.

#### Goal of BWA-MEME

Our goal is to build a practical and efficient alignment software that leverages learned index and suffix array search in the seeding phase. The accuracy of the short read alignment is important, therefore the output must be identical with that of BWA-MEM2. This paper considers running alignment software on CPUs only. However, we believe it can be further accelerated by using hardware acceleration.

## 4 Design of BWA-MEME

### 4.1 P-RMI: Partially-3-layer RMI

We present P-RMI that enables efficient suffix array search by making two enhancements to 2-layer RMI:

#### Mitigating data imbalance

Due to redundant sequences in the reference DNA, using naïve 2 or 3 layer RMI results in imbalanced distribution of data in the leaf models. The number of keys per leaf model forms a long tail distribution, with a small number of leaf models occupying a significant fraction of keys. A model trained with a small subset of data generally has a higher prediction accuracy. Therefore, mitigating data imbalance is necessary to improve the prediction accuracy of RMI. For the human reference DNA and a 2-layer RMI constructed with 2^28^ leaf models, 85% of the leaf models have data, and the rest are empty. Among the non-empty leaves, 0.22% of them hold 16.5% of key-value data from the suffix array.

To mitigate the imbalance in data distribution with minimum overhead, we introduce partially-3-layer RMI (P-RMI). Instead of fully employing the third layer models, P-RMI adaptively adds an additional layer of models only to the second layer models which are suffering from elongated lookup time due to the imbalance in data distribution. Training P-RMI is done layer by layer which is the same as RMI. If the number of assigned keys in the second layer model exceeds the number of key thresholds, an additional layer of models is added to the second layer model. Accordingly, the prediction of the second layer model shifts its role to assign keys to the models in the added layer. The number of models in the additional layer is calculated according to the number of keys assigned and the target average keys per leaf model. When an additional layer is added, an indicator bit is set and stored with the parameters.

#### Defining the error bound for last-mile search

To minimize the number of memory access in the last mile search, we choose to perform a binary search within the predefined error bound instead of using the exponential search or linear search. However, RMI does not guarantee an error bound for the keys that are not provided in the training phase. The original RMI work provides a solution that can be used in 2-layer RMI when all models within the RMI are monotonic (Kraska et al., 2018; Marcus et al., 2020; Rashelbach et al., 2020). However, it does not generalize for 3 layer RMI even if the models are monotonic. Thus, we extend the solution to work on our P-RMI.

Our main observation is that the error bound can be guaranteed for the multi-layer RMI, if the leaf models are monotonically increasing function and the index of the leaf model that a key is assigned to is monotonic with regard to the input key. We omit the proof here. Therefore, we build P-RMI with two design constraints. First, P-RMI forces models to be a monotonically increasing function. Second, the P-RMI constrains that the models form a tree. This makes the index of the leaf model monotonically increasing with regard to the input key. ML models used in the leaf models of P-RMI and the position of keys are both monotonically increasing functions of the key. Therefore, a prediction error is also a monotonically increasing function of the key as it is the difference between the output of the leaf models and the position of keys. Hence, finding the minimum and maximum error of all keys that can be assigned to the leaf models guarantees an error bound to be defined for arbitrary input keys, including duplicate keys or keys not seen in the training phase.

#### P-RMI configuration

The following factors determine the performance of a P-RMI: number of models and types of ML models. Using a large number of models results in a smaller error bound but increases the size of P-RMI. To balance between the lookup performance and the size of P-RMI, BWA-MEME choose a number of models in the second and additional layer to match the target average keys per model. Also, the additional layer is added to the second layer models where a number of keys exceed the number of keys threshold. We choose 20 as the target average keys per model and 500 as the number of keys threshold. Another important configuration is the type of model as it affects both the prediction accuracy and the size of the index. We found the best performance can be obtained using bit shift operation in the first layer, linear regression models in the second layer, and linear spline models in the additional layer. For the human reference DNA, P-RMI has 2^28^ models in the second layer and 48,047,097 models in the additional layer resulting in a total of 7.6GB.

### 4.2 Exact match search with P-RMI

BWA-MEME replaces the exact match search algorithm of BWA-MEM2 with an exact match search algorithm (Exact-MEME) based on suffix array search that uses P-RMI.

#### Tokenization of query sequence

Tokenization is a procedure that encodes a variable length suffix or a substring of the short read into a numerical key. The tokenized key should preserve the lexicographical order and be expressive enough to represent the string key. We observe that most suffixes longer than a certain length become unique suffix strings in the reference DNA sequence. Therefore, we choose to use the first *K* characters of the query sequence and apply 2-bit encoding to the characters. One problem that arises is using K larger than 32 makes the computation expensive in the models of RMI (Wang et al., 2020). For example, using a 128-bit (64 bases) key requires a 128-bit machine learning model to be used in RMI. However, no mainstream processors have hardware support for the floating-point operation with bits larger than 64. This results in a simple 128-bit linear regression model to be 5 times or much slower than the 64-bit linear regression model. Also, we observe that *K* larger than 32 has marginal gain in prediction accuracy, thus we select 32 for *K*.

Tokenization of variable-length string is done in two steps. The first 32 characters are obtained from each variable-length string by padding an arbitrary character to the string if the string is shorter than 32 and trimming characters if it is longer. The 32 characters are then encoded into a numeric key using the 2-bit encoding of bases.

#### P-RMI lookup

P-RMI stores an indicator bit in the second layer models that tells an additional layer of models is added or not. If the indicator bit is not set, the prediction and error value from the second layer is chosen as the output. If the indicator bit is set, the start and end indices of the third layer are obtained from the error value. According to the prediction of the second layer model, the leaf model is selected among the third layer models between the start and end indices.

The error obtained from P-RMI includes maximum error and minimum error which defines the error bound. A binary search is performed within the error bound to find the sorted position of the query sequence in the suffix array. If the sorted position is found and the query sequence exactly matches the corresponding suffix, the sorted position is selected as the longest exact match position. If the sorted position of the query sequence is between two sequential suffixes, the position of suffix with the longer exact match is selected. For notation simplicity, we refer to the longest exact match found in Exact-MEME as LEM.

#### Reducing memory accesses in the last mile search

The last mile search stage accounts for a significant portion of memory accesses. Therefore it is important to reduce the memory accesses. During the last mile search, comparisons are performed several times between suffixes and the input substring. To compare a suffix and the input substring, characters of suffix should be obtained from the reference DNA sequence which requires two random memory accesses. The first memory access is made to the suffix array which contains the position of the suffix in the reference DNA sequence. Subsequently, another memory access is made to the reference DNA sequence to retrieve the suffix characters. The two memory access are random and likely to cause CPU cache misses. To reduce the cache misses, BWA-MEME co-locates position value and the characters of the suffix in a single index data-structure. In particular, the position value and the 32 characters (64-bit) of the suffixes are co-located, allowing any substring shorter than 33 can be compared with a single CPU cache miss. For the human reference DNA sequence, using 32-character suffixes results in 49GB of memory usage.

### 4.3 Making SMEM Search Efficient

This section presents how BWA-MEME find SMEMs using the Exact-MEME algorithm.

#### Extension with Exact-MEME algorithm

The extension finds a substring that has the maximum number of hits but is less than or equal to the hit threshold. However, the Exact-MEME algorithm finds only the longest exact match (LEM) position without progressively extending a substring. To remedy this, BWA-MEME designs an extension function that performs a linear search starting from the output LEM position of the Exact-MEME algorithm. The key insight we leverage is that exact matching positions are all sequentially positioned in the suffix array and the exact match length of query in suffixes monotonically decreases as distance increases from the LEM position. The algorithm of the extension function consists of 3 steps. First, the query of extension function is given to the Exact-MEME algorithm which outputs the LEM position of the query in the suffix array. Next, a linear search is performed starting from the LEM position. The linear search defines an exact match range of suffixes where the input substring has a partial exact match. The size of the exact match range must be largest as possible but is less than or equal to the hit threshold. Also, input substring and all suffixes in the exact match range must have an exact match length longer than that of suffixes not in the exact match range. Finally, the start position of the exact match range, the minimum exact match length inside the exact match range, and the size of the exact match range (number of hits) are returned as the output.

#### Reducing redundant extensions without using LEP bits

Reducing the number of redundant extensions in the SMEM search algorithm is necessary to minimize the number of exact match searches. SMEM-BWA and SMEM-ERT achieve this by tracking the change in number of hits during the first forward extension. The LEP bits obtained from the forward extension are used to reduce redundant extensions. However, unlike SMEM-BWA or SMEM-ERT, it is infeasible to track the change in number of hits while extending a substring using the Exact-MEME algorithm. Therefore, we design a new SMEM search algorithm that does not require LEP bits and uses the same or less number of extensions compared to SMEM-BWA or SMEM-ERT. We observe that repeatedly performing backward and forward extensions starting from the pivot point finds all SMEMs without redundant extensions. The extensions are performed until the extension no longer includes the pivot point. We show the correctness of SMEM output by proving the SMEM output is identical with SMEM-ERT. The proof can be found in the Supplementary Material.

### 4.4 P-RMI Lookup Result Reuse

To identify all possible alignment candidates, the seeding algorithm performs multiple SMEM searches on each short read with various pivot points and thresholds. This results in higher sensitivity in alignment (Li, 2013), but also numerous exact match searches of substrings that have long overlap with each other. But, even if the queries have long overlap with each other, different paths are accessed in P-RMI which incurs random memory accesses, which results in CPU cache misses. The P-RMI lookup and last-mile search accounts for most of the computation time in the seeding phase, thus reducing their cost brings further speedup.

#### Substituting P-RMI lookup for ISA lookup

To reduce memory accesses, we use an additional index called inverse suffix array (ISA). At the index building step, BWA-MEME constructs ISA that stores the translation from the reference position to the suffix array position (i.e. *ISA*[*SA*[*j*]] = *j for j* ∈ |*SA*|). Assume a query *Q_i_* where the LEM position in the suffix array is known and a new query *Q_n_* are given. BWA-MEME uses the ISA and LEM position of *Q_i_* to predict the LEM position of *Q_n_* if *Q_i_*[*X* : *X* + *N*] overlaps with *Q_n_*[0 : *N*] where *X* is the start position and N is the length of the overlap. We refer to the overlapping bases between the identified query and the new query as *overlapping bases*. Let SA and idx be the suffix array and LEM position of *Q_i_*, such that *Q_i_* aligns to *SA*[*idx*] position in the reference. Then the overlapping bases must align to *SA*[*idx*] + *X* position in the reference and the LEM position of the overlapping bases in the suffix array is *ISA*[*SA*[*idx*] + *X*]. To find the LEM position of *Qn* where *Qn*[0 : *N*] is the overlapping bases, BWA-MEME performs last-mile search starting at *ISA*[*SA*[*idx*]+*X*] in the suffix array. The benefits of using ISA instead of P-RMI lookup come from better spatial locality and higher prediction accuracy. To predict the LEM position of the *Qn*, ISA is accessed within the length of short read distance from *ISA*[*SA*[*idx*]], whereas P-RMI lookup requires completely random access to memory. Also, long overlaps often lead to unique suffixes which results in better prediction than using P-RMI lookup.

#### When to use lookup result reuse

BWA-MEME decides to reuse P-RMI lookup result in two cases: First, when the short read has a perfect match to the reference DNA sequence, it is guaranteed for the LEM positions of any new queries to align to the perfect exact match position of the short read. The LEM positions of the new queries can be concluded from the LEM position of the perfect exact match without further last-mile search. As a perfect exact match of short read is common in NGS, this reduces a significant amount of memory accesses and CPU cache misses. Second, when the new query partially overlaps with the identified query and the overlapping bases are long enough, the LEM position of the new query is likely to be near the LEM position of the overlapping bases. To find the actual LEM position in the suffix array, we use an exponential search starting from the predicted position.

## 5 Results

We evaluate BWA-MEME to answer the following questions:

- Does BWA-MEME have identical SAM output with BWA-MEM2?
- How does it compare with the state-of-the-art alignment software BWA-MEM2 and ERT?
- How effective is P-RMI compared to original RMIs?
- How does BWA-MEME adapt to various memory sizes in servers?
- How sensitive is BWA-MEME to the sequencing error rates in the short read and mutation ratio of the reference DNA?

### 5.1 Methodology

#### Setup

We ran the experiments on Intel Xeon Gold 5220R @ 2.2 GHz with 24 cores, 32KB L1, 24 MB L2, 35.75 MB L3 caches, running Ubuntu 20.04 (Linux kernel 5.4.0). We used the numactl utility to force all memory allocations to a single socket. Unless noted otherwise, 48 threads were used for all experiments with hyper-threading. To analyze the memory access characteristics, we used the Intel Vtunes profiler.

#### Dataset

We use the reference human genome assembly (human_g1k_v37) and 16 short reads datasets—7 from Illumina Platinum Genomes (Eberle et al., 2017) and 9 from 1,000 Genomes Project Phase 3 (Consortium et al., 2015). Details are included in the Supplementary Material.

#### Implementation

All algorithms of BWA-MEME are implemented in 3.5k lines of C++ code and integrated into BWA-MEM2 code. BWA-MEM2 is the most widely used alignment software with various features used by researchers. To guarantee BWA-MEME supports all features supported in BWA-MEM2, we choose to replace the seeding algorithm in BWA-MEM2 with our seeding algorithm. For the correctness of BWA-MEME, we verified BWA-MEME and BWA-MEM2 have identical SAM outputs in all 16 short read datasets. We implement the training process of P-RMI on top of the open-source code based on Rust from the authors of the learned index (Kraska et al., 2018). The training process outputs the model parameters of P-RMI in binary data which is stored along with the indices of the reference DNA. The model parameters are loaded in the index loading step of BWA-MEME and used for the seeding algorithm.

### 5.2 Comparing BWA-MEME, BWA-MEM2, and ERT

To demonstrate BWA-MEME delivers significant improvement in seeding throughput, we compare BWA-MEME with two state-of-the-art alignment software, BWA-MEM2 and ERT. Note that we cannot compare with LISA and Sapling because they do not provide complete seeding.

#### Seeding throughput comparison

Figure 4 (a) shows the average seeding throughput of BWA-MEM2, ERT, and BWA-MEME for the 16 short read datasets. The seeding throughput of each alignment software in the figure is normalized with respect to the seeding throughput of BWA-MEM2. The error bars represent the standard deviation of the normalized seeding throughput. BWA-MEME achieves average 3.32x speedup over BWA-MEM2 and average 1.72x speedup over ERT. This is because algorithms in BWA-MEME process exact matches in more memory efficient and cache-friendly manner. Due to its efficient design, BWA-MEME completes the job with 4.60x fewer number of instructions, 8.77x fewer memory accesses, 2.21x fewer last-level cache (LLC) misses, and 4.41x less size of data fetched per read, as shown in Figure 4 (b), (c), (d), and (e), respectively.

**Fig. 4.**
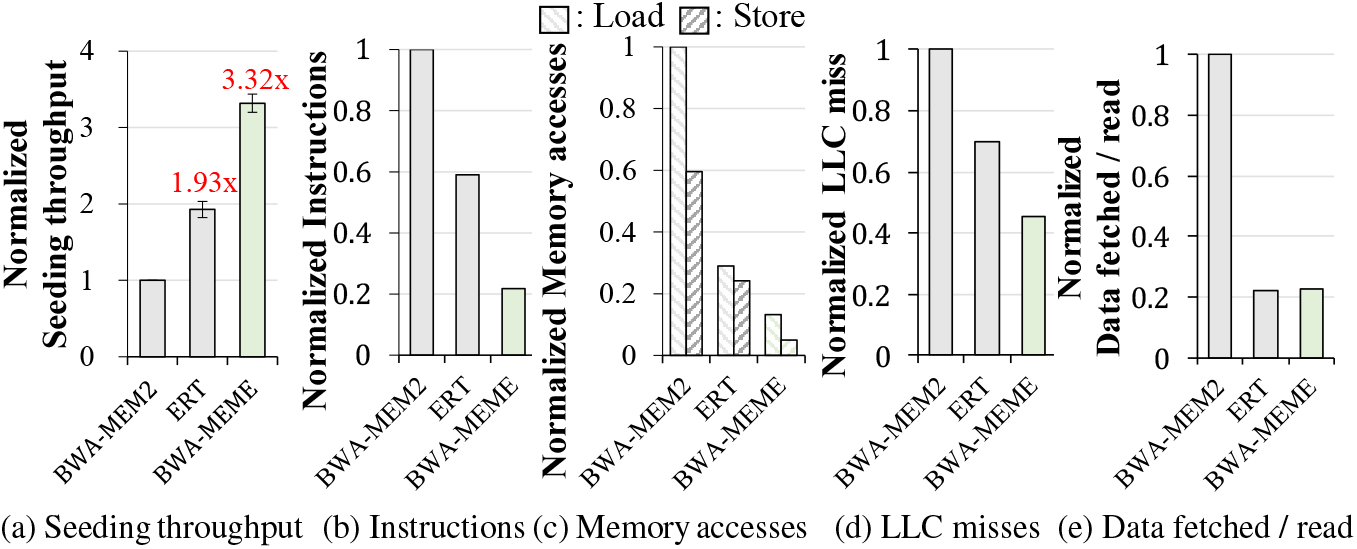
Comparing BWA-MEM2, ERT, and BWA-MEME

#### Implications on alignment throughput

Figure 5 compares the end-to-end alignment throughput of BWA-MEME and memory requirement. BWA-MEME achieves up to 1.42x and 1.12x speedups over BWA-MEM2 and ERT. Note that the seeding phase accounts for average 50% of the runtime in BWA-MEM2. Our seeding algorithm dramatically enhances the seeding throughput. As a result the seeding phase accounts for 29.9% in BWA-MEME as shown in Figure 8.

**Fig. 5.**
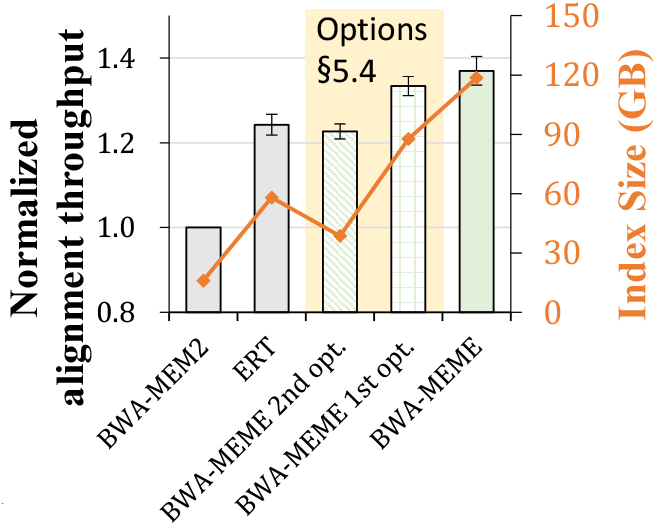
Alignment throughput of BWA-MEM2, ERT, variants of BWA-MEME,and BWA-MEME

### 5.3 Performance Benefit of P-RMI

P-RMI contributes to higher seeding throughput because it provides accurate prediction and a small error bound compared to the original RMIs. Figure 6 (a) and (b) compare the number of last mile search and seeding throughputs of using P-RMI or original RMIs in BWA-MEME. The seeding throughput is normalized by the seeding throughput of BWA-MEM2. For this evaluation, we use the first 10 million short reads in ERR3239276 from 1,000 Genomes Project Phase 3. To accurately measure the effect of choosing the RMI structure, we excluded the acceleration by lookup result reuse. For 2-layer and 3-layer RMI, we used the linear spline model in the leaf models and otherwise linear regression model which showed the best performance in the RMI optimizer (Marcus et al., 2020). In all cases, the number of leaf models is fixed to 2^28^. For a fair comparison of 3-layer RMI and P-RMI, we used 48,047,097 models in the second layer of 3-layer RMI which is the same number of models used in the additional layer of P-RMI. We applied a binary search when an error bound is provided and an exponential search when an error bound is not provided. As shown in Figure 6 (a) and (b), using an exponential search generally requires more number of last mile search compared to using a binary search. The 3-layer RMI makes accurate predictions compared to the 2-layer RMI, however, it incurs more CPU cache misses while inferencing the models and performing the last mile search. Therefore using the 2-layer RMI with error bound outperforms the 3-layer RMI in seeding throughput. P-RMI combines the two advantages of using a binary search within the error bound and higher prediction accuracy of 3-layer RMI. Hence, P-RMI successfully outperforms the existing RMIs in seeding throughput, at most 29.3%.

**Fig. 6.**
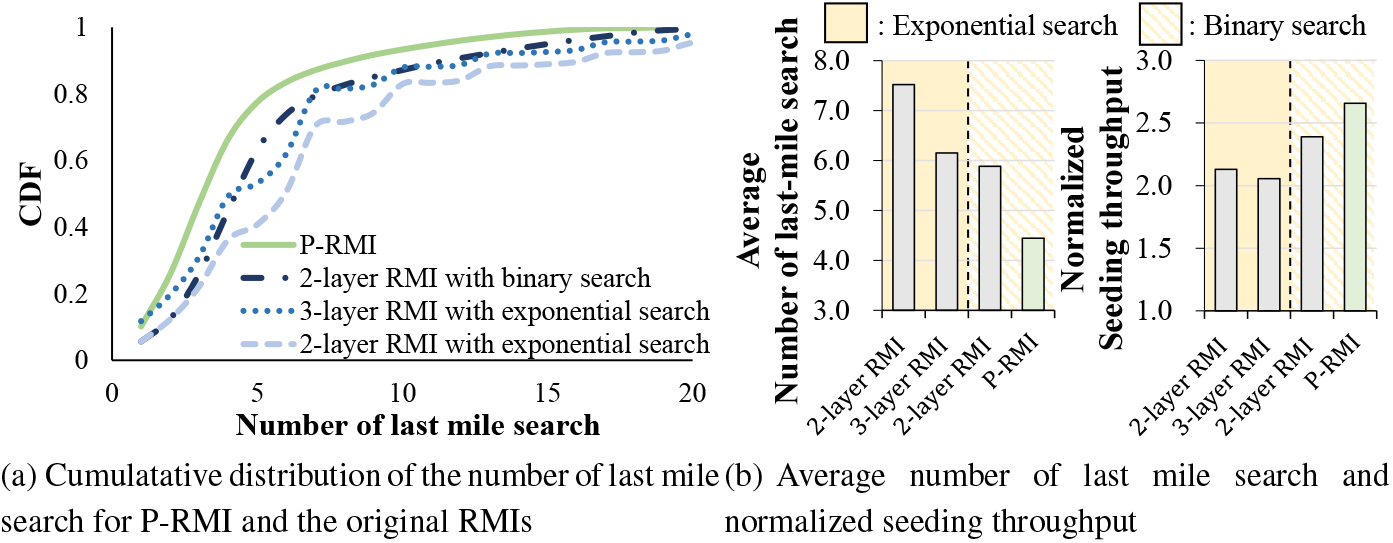
Comparing P-RMI and original RMIs

### 5.4 Memory trade-off design of BWA-MEME

We demonstrate BWA-MEME is able to meet various memory constraints. Figure 5 shows the memory requirement and alignment throughput of each variant of BWA-MEME. BWA-MEME uses 118 GB of memory, which consists of P-RMI, suffix array (SA), 64-bit suffixes (32 characters from §4.2), and inverse suffix array (ISA). BWA-MEME provides two options that selectively load index data-structures to use less memory space, which comes with a tradeoff in throughput. The first option is to exclude ISA used for lookup result reuse, which brings down the memory requirement to 88 GB. The seeding throughput slightly degrades over the full-mode BWA-MEME, but it still outperforms BWA-MEM2 and ERT, achieving 2.69x and 1.33x speedups in seeding and alignment throughput over BWA-MEM2. The second option excludes both the ISA and the 64-bit suffixes, BWA-MEME uses only the P-RMI and suffix array to process seeding. The memory requirement goes down to 38 GB, and BWA-MEME still achieves 1.71x and 1.23x speedups in seeding and alignment throughput over BWA-MEM2, which is similar to those of ERT.

### 5.5 Effect of Mutation and Sequencing Error

BWA-MEME outperforms BWA-MEM2 given various mutation ratios in the reference DNA or sequencing error rate of the short reads. Figure 7 shows the seeding throughput of BWA-MEME in varying sequencing error and mutation ratio. We used 10 million of 200 length synthetic short reads (Li, 2011). The seeding throughput is normalized by that of BWA-MEM2. First, the mutation ratio was fixed to 0.1% to compare the seeding throughput with varying sequencing error rates in the short reads. Next, the sequencing error rate was fixed to 0.1% to compare the seeding throughput with varying mutation ratios of reference. The seeding throughput of BWA-MEME degrades as error increases because the large acceleration comes from processing a long exact matching query with fewer operations. However, BWA-MEME still outperforms BWA-MEM2 in all cases.

**Fig. 7.**
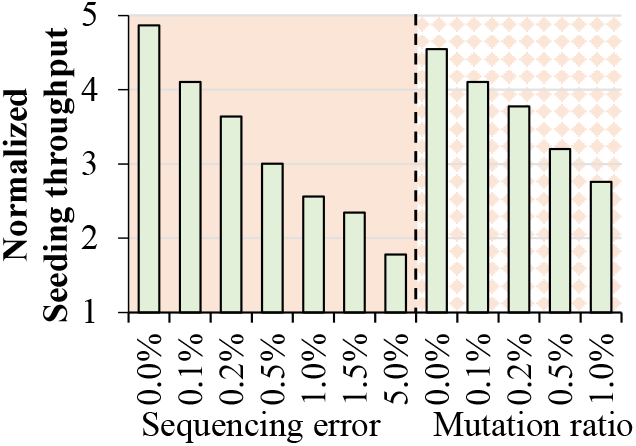
Robustness of BWA-MEME

**Fig. 8.**
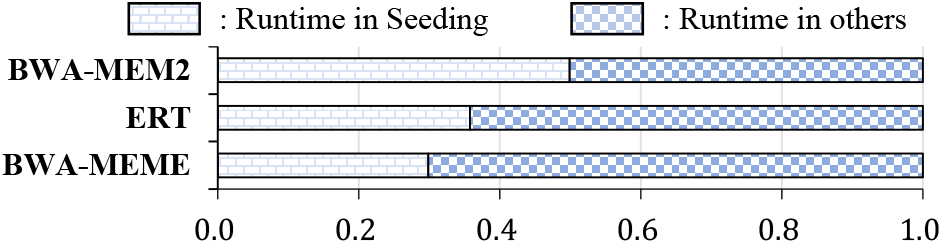
Runtime of the seeding in alignment software

## 6 Discussion

We present BWA-MEME, a new alignment software based on suffix array search and learned index. BWA-MEME introduces novel algorithms and a new learned index structure for seeding algorithm. BWA-MEME replaces the whole seeding algorithm used in BWA-MEM2 while ensuring the identical SAM output with BWA-MEM2. We demonstrated using learned index in suffix array search achieves up to 3.45x and 1.42x speedups in seeding throughput and alignment throughput over BWA-MEM2 by reducing number of instructions 4.60x, memory accesses 8.77x, number of LLC misses 2.21x, and data fetched per read 4.41x. Finally, to accommodate various memory sizes in the server, BWA-MEME provides options to balance the trade-off between seeding throughput and the required memory size.

