## Supplementary Material for "BWA-MEME: BWA-MEM emulated with a machine learning approach"

### EMPLOYING LEARNED INDEX IN THE SUFFIX ARRAY SEARCH REQUIRES $O(1)$ MEMORY ACCESSES WHICH IS INDEPENDENT WITH THE LENGTH OF INPUT SUBSTRING

To find the exact match position of the input substring, the memory accesses incur during the model inference and the last mile search. The number of memory accesses in RMI model inference depends on the number of layers in the RMI and is a constant number. Subsequently, memory accesses occurs in the last mile search where a binary search is performed within the error bound. Each comparison during a binary search requires an $O(1)$ memory accesses. As the error bound is a constant number that is determined at the index building step, the binary search incurs $O(1)$ memory accesses. Both RMI inference and the last mile search incur $O(1)$ memory accesses. Therefore, the exact match search problem can be solved with $O(1)$ memory accesses when employing the learned index in the suffix array search.

### PROOF: SMEM SEARCH OF BWA-MEME HAS IDENTICAL SMEM OUTPUT WITH SMEM-ERT

Let $R$ be a short read sequence that consists of A, C, G, T. Let $R[i, j]$ denote the substring starting at position i and ending at position j of short read $R$. The SMEM searching stage of SMEM-ERT starts at the pivot point. The backward and forward extensions are repeatedly performed until the forward extension reaches the end of the obtained LEP bits. The forward extension starts at the point where the backward extension ends, and the backward extension always starts at the nearest point where the LEP bit is set to 1. We prove performing backward extension and forward extension without obtaining LEP bits has identical SMEM output.

**Theorem 0.1.** *Repeatedly performing backward and forward extensions starting from the pivot point finds all SMEMs that are identical with SMEM-ERT.*

**Definition 0.1** (Extension)**.** Forward extension performed in the point $Pos$ of the short read $R$ is denoted $forward(pos)$. Likewise, backward extension performed in the point $Pos$ of the short read $R$ is denoted $backward(Pos)$. The extension is performed until it can't be further extended and the output of the extension is the position where the extension ends. Hence, $forward(backward(Pos)) \geq Pos$ should always hold.

**Definition 0.2** (Left extension point bit)**.** The left extension point bits are obtained in the forward extension in the pivot point $P$ of the short read $R$. LEP bit is obtained for all $[P, forward(P)]$, if number of hits of substrings $R[P, P+n]$ and $R[P, P+n+1]$ are different, LEP bit in $P+n$ is set to 1 or otherwise LEP bit is set to 0.

Performing backward and forward extension at the point where the LEP bit is set to 1 is the same algorithm with SMEM-ERT. Thus we prove performing backward and forward extension in the point where the LEP bit is set to 0 have an identical result with starting from the closest point which have LEP bit set to 1. The proof of Theorem 0.1 directly follows from the next three lemmas

**Lemma 0.2.** *Let pos be a point where LEP bit is set to 0 and P a pivot point of the short read. For* $\forall p_s \in [backward(pos), pos]$, $forward(p_s) > pos$.

*Proof.* From extension definition, $forward(p_s)$ should be equal or larger than $pos$. If $forward(p_s) \equiv pos$, there exists unique hits of substring $R[p_s, pos]$ in reference. For $\forall p \in [p_s, pos]$, there exists hits of substring $R[P, pos]$ exists where $R[P, pos+1]$ does not exact match. It is contradiction to assumption that LEP bit is set to 0 in $pos$, therefore for $\forall p_s \in [backward(pos), pos]$, $forward(p_s) > pos$. □

**Lemma 0.3.** *Let pos be a position where LEP bit is set to 0 in the SMEM searching stage. For all pos,* $backward(pos) \equiv backward(pos+1)$.

*Proof.* We divide the possible cases of $backward(pos)$ and $backward(pos + 1)$ in to three.

- Case 1 $backward(pos) > backward(pos + 1)$: Backward extension from $pos$ cannot be shorter than backward extension from $pos + 1$ which leads to a contradiction with Definition 0.2.
- Case 2 $backward(pos) < backward(pos + 1])$: $backward(pos)$ can be smaller than $backward(pos + 1])$ only if $forward(backward(pos)) \equiv pos$. If $forward(backward(pos)) > pos$, it is contradiction to $backward(pos) < backward(pos + 1])$. Also, from lemma 0.2 $forward(backward(pos)) \equiv pos$ is contradiction to the assumption that LEP bit is set to 0 in $pos$.
- Case 3 $backward(pos) = backward(pos + 1])$: As case 1 and case 2 are excluded, for all $pos$, $backward(pos)$ and $backward(pos + 1])$ should be identical.

All cases except case 3 lead to contradiction therefore we have that $backward(pos) \equiv backward(pos + 1])$ for any position where LEP bit is set to 0. □

**Lemma 0.4.** *Let $pos_1$ be the closest position from pos where LEP bit is set to 1 in forward direction. For all $p \in [pos, pos_1], \exists C \in [0, pos)$, s.t. $C = backward(p)$. Thus, performing backward extension and forward extension sequentially from $\forall p \in [pos, pos_1]$ results in $forward(C)$.*

*Proof.* From Lemma 0.3, it is given $backward(pos) \equiv backward(pos + 1)$. For $\forall n \in [1, pos_1 - pos)$, LEP bit in position $pos + n$ is 0 and it is proven from Lemma 0.3 that $backward(pos + n) = backward(pos + n + 1)$. Therefore, for $\forall p \in [pos, pos_1], \exists C \in [0, pos)$, s.t. $C = backward(p)$ and $forward(C) \equiv forward(backward(p)])$

□

### SHORT READ DATASET USED IN EXPERIMENT

| Dataset | read length | number of reads | Source |
|---|---|---|---|
| ERR194146_1 | 101 | 813180578 | Illumina Platinum Genomes |
| ERR194147_1 | 101 | 787265109 | Illumina Platinum Genomes |
| ERR194158_1 | 101 | 859371011 | Illumina Platinum Genomes |
| ERR194159_1 | 101 | 707646124 | Illumina Platinum Genomes |
| ERR194160_1 | 101 | 775617169 | Illumina Platinum Genomes |
| ERR194161_1 | 101 | 843454257 | Illumina Platinum Genomes |
| ERR3239276_1 | 150 | 396570406 | 1000 Genomes Project Phase 3 |
| ERR3239277_1 | 150 | 363937308 | 1000 Genomes Project Phase 3 |
| ERR3239278_1 | 150 | 342631544 | 1000 Genomes Project Phase 3 |
| ERR3239279_1 | 150 | 420210145 | 1000 Genomes Project Phase 3 |
| ERR3239280_1 | 150 | 391766960 | 1000 Genomes Project Phase 3 |
| ERR3239281_1 | 150 | 365635559 | 1000 Genomes Project Phase 3 |
| ERR3239282_1 | 150 | 367637337 | 1000 Genomes Project Phase 3 |
| ERR3239283_1 | 150 | 391766960 | 1000 Genomes Project Phase 3 |
| ERR3239284_1 | 150 | 374824132 | 1000 Genomes Project Phase 3 |

**Table S1.** Short read data used for evalution.

### COMPARISON OF SUPPORTED FEATURES

| Seeding algorithm | Exact match search of arbitrary substring | Seeding stage 1 | Seeding stage 2 | Seeding stage 3 | Can process full alignment |
|---|---|---|---|---|---|
| Sapling | X | X | X | X | X |
| LISA | O | O | X | X | X |
| BWA-MEME | O | O | O | O | O |
| BWA-MEM2 | O | O | O | O | O |
| ERT | O | O | O | O | O |

**Table S2.** Comparison of algorithm features.

To the best of our knowledge, there are no seeding algorithms based on learned index that have full support for the required functionality of alignment software. Thus BWA-MEME is the first alignment software that leverages learned index in the seeding algorithm. The seeding algorithm of BWA-MEM2 consists of 3 stages. The first stage generates seeds, the second stage reseeds the seeds generated in the first stage, and the last stage generates seeds that have more hits to the reference DNA sequence. The SMEM search in the first stage starts from the start of the short read and is performed until the end of the short read. However, the second and third stage is different from the first stage. The second stage executes the SMEM search algorithm in the middle of the seeds found in the first stage. The third stage performs only the forward extension to find seeds with a hit threshold set to 20. We describe the features supported in each algorithm in Table S2

### PSEUDO ALGORITHMS OF BWA-MEME

**Algorithm S1.** Tokenization of query sequence

```
Input: Query sequence Q
Output: Tokenized sequence
procedure GET_KEY_OF_READ(Q)
    Token = 0
    for i ← 0 to min(len(Q),32) do
        c = Q[i]
        if c == Ambiguous base then
            Break
        Token = Token << 2
        Token = Token | 2bit_encode(c)
    while i < 32 do
        Token = Token << 2
    Return Token
```

**Algorithm S2.** Lookup of P-RMI

```
Input: Tokenized query sequence
Output: Predicted position and Error of tokenized query sequence
procedure P-RMI_LOOKUP(Token)
    model_index ← first_layer_model(Token)
    Pred, Err ← second_layer_models(Token, model_index)
    if Err >> 63 then
        Third_model_start_index ← (Err >> 32) & 0x7fffffff
        Third_model_max_index ← Err & 0xffffffff
        Pred = min(Pred, Third_model_max_index)
        model_index = Third_model_start_index + Pred
        Pred, Err = Additional_layer_models(Token, model_index )
    Return Pred, Err                    ▷ Err contains Lower_error and Upper_error
```

**Algorithm S3.** Calculating upper error and lower error of leaf model N

```
Input: (Key, position) dataset K[N] of N leaf models and model n
Output: Upper_Error and Lower_Error of model n
procedure ErrorRangeCal(K[N],n)
    Keys ← K[n-1, -1]+K[n, :]+K[n+1, 0]
    for (Token, Position) ∈ Keys do
        Pred = rmi_lookup(Token)
        Error = Position − Pred
        if Upper_Error < Error then
            Upper_Error = Error
        if Lower_Error > Error then
            Lower_Error = Error
    Lower_error = abs(Lower_error)                ▷ absolute value of Lower error
    Err = Lower_error « 32 | Upper_error
    Return Err
```

**Algorithm S4.** Exact-MEME algorithm

```
Input: Query sequence Q
Output: MEM position and length of MEM
procedure LEM_SEARCH(Q)
    Token ← Tokenization(Q)
    Pred,Err ← P-RMI_lookup(Token)
    Lower_error ← (Err >> 32) & 0x7fffffff
    Upper_error ← Err & 0xffffffff
    Search_bound ← { Pred - Lower_error, Pred + Upper_error }
    MEM position, Length ← BinarySearch(Q, Search_bound)
    Return MEM position, Length
```

**Algorithm S5.** Extension using Exact-MEME algorithm

```
Input: Query sequence Q and the hit_threshold
Output: start position of hit range, length of MEM, number of hits
procedure COMPARE(Q, pos)
    return exact match length of Q and SA[pos]
procedure EXTENSION(Q, hit_threshold)
    Lem_pos, Lem_len ← Lem_search(Q)
    upper_b = Lem_pos + 1; lower_b = Lem_pos - 1;
    while upper_b - lower_b - 1 < hit_threshold do
        while Lem_len == compare(Q, upper_b) do
            upper_b = upper_b + 1
        while Lem_len == compare(Q, lower_b) do
            lower_b = lower_b - 1
        last_lem_len = Lem_len
        Lem_len = max(compare(Q, upper_b), compare(Q, lower_b))
    return (lower_b+1), last_lem_len, (upper_b-lower_b-1)
```

**Algorithm S6.** SMEM-MEME algorithm for BWA-MEME

```
Input: short read, pivot point, hit_threshold, min_seed_len
Output: SMEM list
procedure SMEM_SEARCH(read, pivot, hit_threshold, min_seed_len)
    smem_list ← []; read_rc ← read.reverse_complement();
    search_pivot = pivot
    while 1 do
        Q = read_rc[len(read)-search_pivot: -1]                ▷ Backward extension
        _, len, _ ← extension(Q, hit_threshold)
        search_pivot = search_pivot - len + 1
        if search_pivot > pivot then
            break                      ▷ Extension no longer include pivot point
        Q = read[search_pivot:-1]                               ▷ Forward extension
        pos, len, num ← extension(Q, hit_threshold)
        if num < hit_threshold and len > min_seed_len then
            for Ref_Pos in SA[pos,...,pos + num] do
                smem_list.append( Ref_Pos )
        search_pivot = search_pivot + len
    Return smem_list
```
